# The fitness landscape of the codon space across environments

**DOI:** 10.1101/252395

**Authors:** Inès Fragata, Sebastian Matuszewski, Mark A. Schmitz, Thomas Bataillon, Jeffrey D. Jensen, Claudia Bank

## Abstract

Fitness landscapes map the relationship between genotypes and fitness. However, most fitness landscape studies ignore the genetic architecture imposed by the codon table and thereby neglect the potential role of synonymous mutations. To quantify the fitness effects of synonymous mutations and their potential impact on adaptation on a fitness landscape, we use a new software based on Bayesian Monte Carlo Markov Chain methods and reestimate selection coefficients of all possible codon mutations across 9 amino-acid positions in *Saccharomyces cerevisiae* Hsp90 across 6 environments. We quantify the distribution of fitness effects of synonymous mutations and show that it is dominated by many mutations of small or no effect and few mutations of larger effect. We then compare the shape of the codon fitness landscape across amino-acid positions and environments, and quantify how the consideration of synonymous fitness effects changes the evolutionary dynamics on these fitness landscapes. Together these results highlight a possible role of synonymous mutations in adaptation and indicate the potential mis-inference when they are neglected in fitness landscape studies.

## 1 Introduction

By considering the relationship between genotype and fitness as a topographic map, Wright (1932) created the concept of a fitness landscape. During the last century this concept has been adopted across various subfields of the sciences, and it has been used extensively to study how populations may adapt to novel environments (Perfeito *et al.*, 2011; De Visser and Krug, 2014; Gorter *et al.*, 2018). Only recently have technological and experimental advances enabled the assessment of large empirical fitness landscapes at high resolution (Weinreich *et al.*, 2006; Hietpas *et al.*, 2013; Bank *et al.*, 2014; Wu *et al.*, 2016; Bank *et al.*, 2016). Wright (1932) noted early on that a complete fitness landscape with *L* loci, each of which has *k* alleles, results in a hypercube of *k^L^* genotypes. This enormous dimensionality can never be fully sampled and therefore enforces a careful and limited choice of the mutations that may be assayed in any given experiment. Thus, most fitness landscape studies to date have only considered amino-acid changing mutations (e.g. Bank *et al.*, 2016; Wu *et al.*, 2016). Only considering the genotype-fitness relationship at the amino-acid level entails the risk of misrepresenting the true underlying fitness landscape and, thus, the potential routes along which adaptive walks may proceed.

Firstly, mutations in an amino-acid based fitness landscape are, by definition, non-synonymous. This neglects the accumulating evidence from both comparative and experimental studies that synonymous mutations (i.e., mutations that change the codon but not the encoded amino-acid) can display non-negligible fitness effects (Singh *et al.*, 2007; Drummond and Wilke, 2008; Kudla *et al.*, 2009; Zhou *et al.*, 2009; Lind *et al.*, 2010; Plotkin and Kudla, 2011; Sauna and Kimchi-Sarfaty, 2011; Agashe *et al.*, 2013; Bailey *et al.*, 2014; Firnberg *et al.*, 2014; Hunt *et al.*, 2014; Bali and Bebok, 2015; Presnyak *et al.*, 2015; Agashe *et al.*, 2016; Choi and Aquadro, 2016; Knöppel *et al.*, 2016). For example, recent studies have shown that synonymous mutations can affect the speed and accuracy of translation (Drummond and Wilke, 2008; Saunders and Deane, 2010; Plotkin and Kudla, 2011; Bali and Bebok, 2015), mRNA structure (Shabalina *et al.*, 2013; O’Brien *et al.*, 2014; Presnyak *et al.*, 2015), expression in response to environmental changes (Shabalina *et al.*, 2013), and that they are associated with several organismal malfunctions (Parmley and Hurst, 2007; Hunt *et al.*, 2014). Although synonymous effects undoubtedly exist, effect sizes are often small, which has made a systematic characterization difficult. In particular, to our knowledge there exists no study to date that has characterized whether fitness effects of synonymous mutations vary across environments; a finding that could be in concordance with the costs of adaptation that are frequently reported for amino-acid changing mutations (e.g. Bataillon *et al.*, 2011; Wenger *et al.*, 2011; Hietpas *et al.*, 2013; Rodriguez-Verdugo *et al.*, 2014).

Secondly, the consideration of a fitness landscape at the codon level introduces a lower connectivity of the genotypes, i.e. a different topology of the fitness landscape. Whereas from the amino-acid view of the landscape, any amino-acid transition is possible in a single mutational step, a codon-based landscape requires up to three mutational steps to transition from one amino acid to another. (c.f. Fig. 1A). Hence, even a single amino-acid position in the genome contains a fitness landscape that consists of the (4nucleotides)^3loci^ = 64 codons at that position.

**Figure 1:**
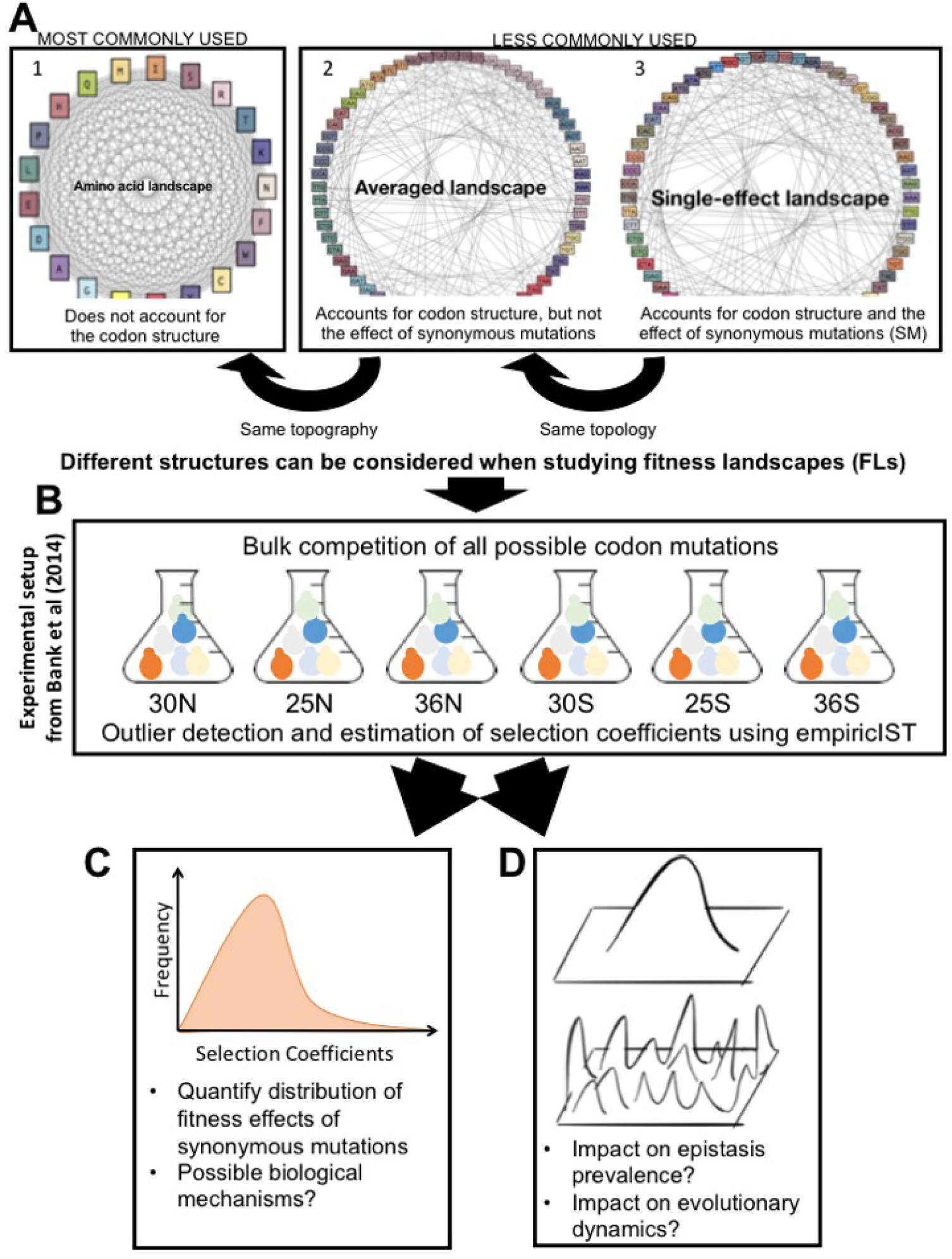
Graphical summary of the study. A) The consideration of the codon structure and the fitness effects of synonymous mutations results in fitness landscapes with different topologies and topographies. The graphs illustrate fitness landscapes at a single amino-acid position. Gray lines indicate single-step mutations and colors indicate potential fitness differences. 1) Many studies implicitly assume that all amino-acids are connected by a single mutational step. 2) The codon table restricts the number of possible substitutions at the amino-acid level and thus results in a different topology. We denote the fitness landscape that accounts for the codon table but neglects the potential effects of synonymous mutations as the averaged landscape (codons that code for the same amino-acid are presented in similar colors). 3) We denote the fitness landscape that considers the individual effect of each codon as the single-effect landscape (each codon has a specific color). B) For this study, we obtained deep mutational scanning data of 54 codon fitness landscapes from (Bank *et al.*, 2014). We infer individual selection coefficients using a newly developed analysis software, empiricIST. C) We quantify the distribution of synonymous fitness effects and perform regression methods to relate these effects to biological mechanisms. D) We quantify the shape of the codon fitness landscapes across environments. We illustrate the consequences of ignoring synonymous effects on the evolutionary dynamics on the landscapes.

We illustrate two aspects of the differences between the amino-acid and codon levels in Fig. 1A. Considering the codon level results in a different topology of the fitness landscape (A1 to A2), and considering effects of synonymous mutations results in a potentially different topography of this codon fitness landscape (A2 to A3). As highlighted by Zagorski *et al.* (2016), a change in the topology of a fitness landscapes can result in dramatically different accessibility of fitness peaks, and the topography further amplifies this effect. For example, a single-nucleotide mutation in a codon-based landscape can result in only 5 to 7 amino-acid changes rather than the 20 total possible amino-acid changes. Thus, at a single amino-acid position, a codon-based fitness landscape (with 64 genotypes) can have multiple local fitness peaks, whereas the corresponding amino-acid landscape (with 21 genotypes) is by definition single-peaked.

Here we quantify the effects of synonymous mutations (Fig. 1C) and study how including synonymous effects modifies the evolutionary dynamics on codon fitness landscapes (Fig. 1D). To this end, we use published data (Bank *et al.*, 2014) from deep mutational scanning (Hietpas *et al.*, 2012; Fowler and Fields, 2014), which consist of codon fitness landscapes of the same 9 amino-acid positions across 6 environments. Our results indicate that the distribution of synonymous effect sizes is heavy-tailed, with many mutations of little effect and a few larger-effect mutations. Furthermore, we compare the shape of the codon fitness landscapes with and without consideration of effects of synonymous mutations. We find that the evolutionary dynamics on these landscapes differ greatly between the two types of landscapes, as local optima created by synonymous effects can stall the progression towards the global optimum of the fitness landscape. Thus, our work calls for a more careful consideration of synonymous effects in future studies of fitness landscapes and adaptive walks.

## 2 Material & Methods

### 2.1 MCMC Method

We implemented a software to infer selection coefficients from deep mutational scanning experiments. The *empiricIST* software is based on a previously developed Bayesian Markov chain Monte Carlo (MCMC) approach (Bank *et al.*, 2014), and is a user-friendly and accurate software for improved growth rate estimation from time-sampled deep-sequencing data. We took advantage of the high accuracy provided by this method to estimate selection coefficients of synonymous mutations.

empiricIST is a software package for 1) processing sequencing count data from deep mutational scanning experiments, 2) estimating growth rates using a Bayesian MCMC approach described in detail in (Bank *et al.*, 2014), and 3) post-processing of growth rate estimates to estimate the shape of the beneficial tail of the distribution of fitness effects (DFE). A detailed description of the software, its usage, and options can be found in the accompanying manual (https://github.com/Matu2083/empiricIST). In the following, we give a brief description of the assumed experimental setup and the model underlying the MCMC and estimation procedure, and by means of simulations compare the accuracy of the results to that obtained from conventional linear regression (Matuszewski *et al.*, 2016).

#### Assumptions of the model and input data

We consider an experiment assessing the fitness of K mutants, labelled *i* ∈ {1,…, *K*}. Each mutant *i* is assumed to be present at initial population size *c*_*i*_ and to grow exponentially at constant rate *r*_*i*_, such that its true abundance at time *t, N*_*i*_(*t*) is given by 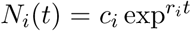.At each sampling time point *t* ∈ {1, …, *T*}, sequencing reads *n*_*i,t*_ are drawn from a multinomial distribution with parameters 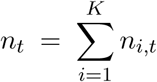 (i.e., the total number of sequencing reads) *p_t_* = (*p_1,t_*, … *p_K,t_*), where 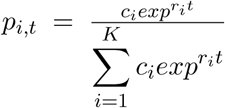 is the relative frequency of mutant *i* in the population at time *t*. Here, time is measured in hours to make results comparable across different environmental conditions (Chevin, 2011; Bank *et al.*, 2014). The software allows for input of either generation or standard time. We furthermore assume that sampling points are independent such that the overall likelihood can be written as the product of the individual likelihoods of each sampling point.

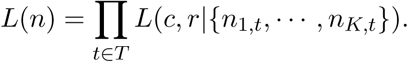

All initial population sizes *c*_*i*_ and growth rates *r*_*i*_ are estimated relative to those of a chosen reference mutant with its initial population size and growth rate arbitrarily set to 10 000 and 1, respectively. Here, the wild-type sequence in laboratory conditions of 30°*C* was used as the reference.

#### MCMC model

We implemented a Metropolis-Hastings algorithm in C++ using flat priors allowing all attainable values *r*_*i*_ ∈ *R*^+^ and *c*_*i*_ ∈ *N* to be realized with equal probability. During the burn-in period the variance of both proposal distributions was adjusted such that the targeted acceptance ratio is around 25%, which optimizes the performance of the MCMC chain (Gelman *et al.*, 1996).

The updated variance of the proposal distribution was calculated using

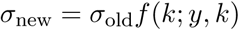

with

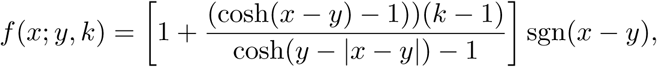

where *x* denotes the targeted acceptance ratio, *y* is the current acceptance ratio, and *k* is a (fixed) scale parameter that restricts the maximal change in the variance of the proposal distribution (Roberts *et al.*, 2001). After discarding the first 100 000 accepted samples (i.e., after the burn-in period), the MCMC was run for an additional 10 000 000 accepted samples. Only every 1000th sample was retained for further analyses, such that the posterior distribution of each parameter was characterized by 10 000 samples overall.

Convergence and mixing were checked by visual inspection of the resulting trace files for all estimated parameters, and by calculating the effective sample sizes (i.e., the number of independent samples) and the Hellinger distance (Boone *et al.*, 2014) between sets of 1000 batched recorded samples. Effective sample sizes were generally larger than 1000 for all parameters, and Hellinger distances below 0.1 indicated convergence and good mixing. To facilitate estimation, we took advantage of the fact that the multinomial distribution is preserved when a subset of the counting variables are observed. This enabled us to split the data set into subsets with 10 mutants each (implicitly treating the other mutants’ sequencing reads as observed). More options such as outlier detection, data imputation, DFE tail-shape estimation are detailed in the supporting Information.

#### Assessing accuracy of the MCMC

To assess the accuracy of the Bayesian MCMC approach, we compared its parameter estimates to those obtained using ordinary least squares (OLS) linear regression of the log-ratios against the number of sequencing reads *n_i,t_* over the different sampling time points (Matuszewski *et al.*, 2016). For that we simulated time-sampled deep sequencing data (implemented in C++; available from https://github.com/Matu2083/empiricIST), assuming that individual mutant growth rates and initial population sizes for each of the *K* mutants are drawn independently from a normal distribution (i.e., *r*_*i*_ ~ 𝒩(1, 0.01)) and a log-normal distribution (i.e., *c*_*i*_ ~ 10 ^𝒩(4,0^-^25)^), respectively. Without loss of generality, we denote the wild-type reference (or any other reference genotype) by *i* = 1 and set its growth rate to 1.

Sequencing reads were then drawn independently for each of the *T* equally spaced time points from a multinomial distribution with parameters *n*_*t*_ (i.e., the number of total sequencing reads per time point) and *p*_*t*_ =(*p*_1_,*t*,…,*p_K_,t* To check the robustness of these results when applied to the real experimental data, we furthermore drew growth rates from a mixture distribution

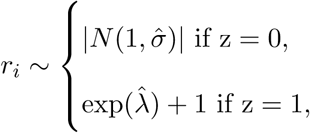

where 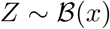 is a Bernoulli-distributed random variable that indicates whether growth rates are drawn from the deleterious part of the DFE (i.e., if *z* = 0) or from the exponential beneficial tail (i.e., if *z* = 1). The parameters 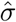, 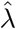, and 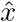 are estimated from the underlying experimental data, and based on growth rate estimates obtained from OLS linear regression.

Finally, the accuracy of the parameter estimates was assessed by computing the mean square error (MSE)

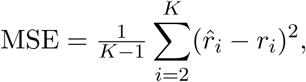

the length of the credibility interval (CI, calculated from the MCMC posterior distribution), and the frequency of the true growth rate lying in the 95% confidence interval obtained via OLS calculated over 100 simulated data sets.

### 2.2 Bayesian MCMC outperforms linear regression

Validating the method with various types of simulated data mimicking the experimental data (as detailed in the supplementary Material) shows that our MCMC generally outperforms ordinary least square regression (OLS). Figure. 2 and S1 show the simulation results. Although the mean square error (MSE) of the MCMC is comparable to that of the OLS when analyzing few time points (i.e., 3 to 5 time points), the MSE of the MCMC decreases faster as the number of time points increases (Fig. 2A).

**Figure 2:**
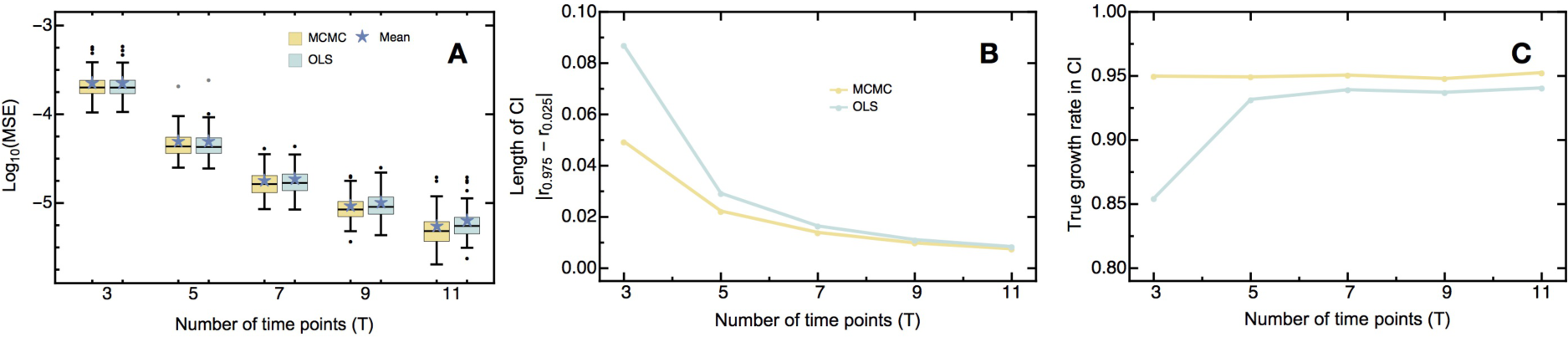
Comparison between performance of *empiricIST* and ordinary least square regression with varying number of time points sampled. We display A) mean square error (MSE), B) size of the credibility interval (CI), and C) the proportion the true growth rate contained in the CI. As shown, *empiricIST* yields an equal or lower MSE than OLS regression, particularly as the number of sampled time points increases. Furthermore, *empiricIST* outperforms the OLS regression regarding the size of the CI and at capturing the true growth rate, even when sampling a small number of time points.

Furthermore, when analyzing few time points, the length of the credibility interval (CI) is significantly smaller for the MCMC than the corresponding confidence interval of the OLS regression (Fig 2B). While the difference between the length of the confidence intervals decreases as the number of time samples *T* increases, the size of the CI from the MCMC always remains smaller, which implies that it yields more precise and accurate results than the conventional OLS regression. Most importantly, and unlike the OLS regression, the CI of the MCMC remains well calibrated along the entire range of parameters that were tested (cf. Fig. 2 C for illustration across a range of time points), despite being generally narrower than its OLS counterpart.

Apart from its main program - the Bayesian MCMC program - *empiricIST* provides Python and shell scripts for data pre- and post-processing. Details about their usage and options are given in the accompanying manual. Here we outline the two different options that are available for dealing with outliers in the sequencing data - i.e., outlier detection and data imputation - and explain the DFE tail-shape estimation.

#### *Outlier detection in* empiricIST

As an alternative to treating outliers as unobserved (i.e., as missing data), we also implemented an approach in which data points identified as outliers were imputed (see SI). For that we again used the linear regression of the log ratios of the mutant’s read number to the total number of reads at each individual time point (i.e., the ‘total’ normalization, *sensu* Bank *et al.*, 2014), and classified as outliers data points that exceed the DFBETA cutoff of 2 and that had an absolute studentized residual bigger than 3. In comparison to other reasonable and established outlier criteria, this approach proved to be more cautious as exemplified by the higher specificity and lower sensitivity (Fig. 2, Fig. S1). By combining two independent outlier criteria (i.e., the DFBETA statistic and the studentized residuals), this approach ensures that data points identified as outliers have leverage effects (i.e., change the slope considerably) and are in conflict (meaning that are very different in comparison) with the remaining data points. Thus, to minimize changes in the original experimental data we took an extremely conservative approach, such that only those data points that stand out as extreme outliers will be imputed.

When comparing the mean square error (MSE) over 100 simulated data sets across different outlier detection methods, we find that the MSE increases with the proportion of outliers in the data set, independent of the method used. Imputing data points generally improves the accuracy of the parameter estimates compared to treating outliers as missing data (Bank *et al.*, 2014,; Fig. S2, S3). Expectedly, when there are no outliers in the data, normalization to the wild type displays the lowest error (c.f. Bank *et al.*, 2014; Matuszewski *et al.*, 2016). However, with only 1% outliers in the data, the error of the normalization to the wild type is comparable to that of the normalization to the total number of reads and performs increasingly worse as the proportion of outliers in the data increases (Fig. S2). Note that in the presence of outliers, using any outlier method improves growth rate estimates considerably.

#### *Estimating the shape for the beneficial tail with* empiricIST

Finally, *empiricIST* contains a Python script for estimating the shape of the beneficial tail of the DFE. It is often believed that these effects typically follow an exponential distribution (Gillespie, 1983, 1984) characterized by many small, nearly-neutral mutations and a few strongly beneficial mutations. Using extreme value theory, it is possible to test whether experimental data complies with that assumption (and falls into the Gumbel domain), or whether the data is better represented by distributions from the Weibull domain (i.e., bounded distributions that decay more rapidly than an exponential distribution, implying more small-effect mutations) or from the Frechet domain (i.e., distributions decaying less rapidly than an exponential distribution implying an excess of large-effect mutations; see also Beisel *et al.*, 2007). Additional information about the different types of distributions and likelihood estimation are available in the section on DFE estimation in the SI (SI-Additional Material and Methods, section DFE tail-shape estimation, Fig S4).

We analyzed the power of the maximum-likelihood method to make this distinction by simulating 1000 Generalized Pareto Distribution (GPD) data sets for different underlying shape parameter (*κ*) values (spanning across all three GDP domains) and varying sample sizes. We find that for small sample sizes (Fig. S4A, B) 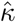 displays a large variance and a slight negative bias, in particular, if the underlying shape parameter is from the Weibull domain (i.e., *κ <* 0). This bias is caused by a (numerical) discontinuity in the log-likelihood function around *κ*= −1 (eq. S3 in SI), causing *κ* to consistently deviate (Rokyta *et al.*, 2008). As sample size increases, however, the variance of the maximum-likelihood estimate decreases and its bias vanishes (Fig. S4C, D). Furthermore, while *κ* typically falls into the correct domain (even for low sample sizes), the statistical power for detecting deviations from the null hypothesis (i.e., whether *H*_0_: *κ* = 0) is low (unless sample sizes are large).

### 2.3 Experimental data

The data used in this study were originally obtained in Bank *et al.* (2014) using the EMPIRIC approach (Hietpas *et al.*, 2011, 2012). Briefly, single-codon-substitution libraries were generated using a plasmid constitutively expressing Hsp90. These were then transformed into the *Saccharomyces cerevisiae* DBY288 shutoff strain (Hietpas *et al.*, 2011; Bank *et al.*, 2014) using the lithium acetate method. Amplification occurred initially for 12 hours at 30^°^C in nonselective galactose medium with ampicillin (100*μ*g/ ml, please see details of medium composition in Bank *et al.* (2014)). These were then transferred to selective dextrose medium, also at 30^°^C, to initiate shutoff of the wild type copy of Hsp90. Bulk competition started after 8 hours in this selective medium, under six different environmental conditions (25^°^C, 30^°^C, 36^°^C, 25^°^C + S, 30^°^C + S, and 36^°^C + S, where S represents the addition of 0.5 M sodium chloride). For simplicity, we will refer to these conditions as normal medium or high-salt medium, and abbreviate these by 25N and 25S, for example, when additionally referring to the 25^°^C environment. Samples were taken at several time points during the experiment (Table S1) and stored at −80^°^C for posterior DNA isolation and sequencing. Sequencing was performed by the University of Massachusetts deep sequencing facility, which generated approximately 30 million reads (see also Table S1, Bank *et al.*, 2014). For further details regarding the experimental method please see Hietpas *et al.* (2011, 2012, 2013); Bank *et al.* (2014).

The data set analysed here contains all 576 possible single-codon mutations in a 9-amino-acid region of the C terminal part of Hsp90 (amino acid positions 582 to 590) in *Saccharomyces cerevisiae*. Whereas from most environments only a single replicate was available, we had access to three technical replicates at 30N and two biological replicates at 30S. Populations were originally adapted to the 30N environment. Growth rates for all mutants were estimated using *empiricIST.* Furthermore, to obtain growth rate estimates per amino-acid (residue) position, we pooled nucleotide sequences and jointly estimated growth rates for those nucleotide sequences that resulted in the same amino-acid sequence (see above and SI). All downstream analyses are based on 1000 subsamples of the posterior distribution obtained from *empiricIST*, if not otherwise indicated. Selection coefficients were obtained by normalizing to the median growth rate of all mutations synonymous to the reference sequence as detailed in Bank *et al.* (2014).

#### Distribution of synonymous mutations

We obtained the distribution of synonymous fitness effects across all amino-acid mutations as the difference between the selection coefficient of each individual codon and its corresponding pooled amino-acid estimate. These data were used to perform the analyses in Section *The distribution of synonymous fitness effects.*

#### Quantifying the impact of GC bias

To check whether Illumina sequencing created a GC bias in our data, we estimated the impact of GC content throughout the several steps of data acquisition and selection coefficient estimation. Firstly, because the library composition was not assessed directly for the data sets used in this study, we used the web plot digitizer (https://automeris.io/WebPlotDigitizer/) to obtain the abundance of each of the 64 codons during library construction in the data from Hietpas *et al.* (2011), supplementary Figure 7C in Hietpas *et al.* (2011). We estimated the fraction of each of the 4 nucleotides present and calculated the deviation from the expected 25%. This was done for all three codon positions. We found a positive bias towards AT codons (Fig. S5) in the library construction. To estimate GC bias in the sequencing data obtained after the experiment we calculated how many G or Cs (guanine or cytosine nucleotides) were present in each barcoded codon (minimum 10, maximum 17). A glm (generalized linear model) using the negative binomial family with *Environment* indicated a small positive bias of GC content in the sequence abundance (GC: 0.098, P <0.0001) This bias was also observed when we tested the correlation between CG abundance and selection coefficient for each amino-acid substitution using an ANOVA model including *GC content* and *Environment* (GC: 0.00456, P <0.0001). However, when repeating this analysis using the selection coefficient of synonymous mutations (i.e., after subtracting the amino-acid effect) this bias was no longer significant (GC: 0.00003, P= 0.7244), indicating that the observed GC bias may indeed reflect selection rather than being an artifact of sequencing. Nevertheless, to account for any potential contribution of GC content or its interaction with other mechanisms, we included the GC abundance in the models that were used to identify possible mechanisms causing synonymous fitness effects.

### 2.4 Detecting the effect of synonymous mutations

#### Experimental error and reproducibility of measurements

To assess the reproducibility of measurements, we compared the correlation between selection coefficient estimates across the three 30N and two 30S replicates, and computed the overlap in their growth rate posteriors. For each replicate pair, we calculated the correlation between mutation-specific fitness effects from both the median estimates and 1000 randomly selected posterior samples. The median correlation of fitness effects across pairs of replicates for high salt medium (biological replicates) was 0.84 (lower and upper credibility intervals from 1000 posterior samples: [0.78, 0.88]) and for standard medium (technical replicates) it was 0.98 (lower and upper credibility intervals from 1000 posterior samples: [0.97, 0.99]), confirming that the experimental protocol has an excellent resolution for measuring selection coefficients. An ANOVA test indicated that experimental error was negligible in comparison to the effect of changing medium (Table S2) and confirmed the previously observed strong costs of adaptation (Hietpas *et al.*, 2013).

To quantify whether the *empiricIST* credibility intervals cover the experimental error appropriately, we estimated the overlap between the 95% credibility intervals of the posterior distribution for all pairs of replicates. We observed a large overlap between pairs of replicates (Fig. S6, normal environment - a) Rep1-2: 98%; b) Rep1-3: 91%; c) Rep2-3: 90%; high salt environment - d) Rep1-2: 90%), indicating that the variance between replicates is indeed mostly covered by variance in the posterior distribution, and that we can use *empiricIST* credibility intervals as confidence levels in our analysis.

We used linear models to quantify the contribution of various factors to the estimated effects of synonymous mutations. Model variable names are highlighted throughout the paper using Italics. The respective analyses were performed on the distribution of synonymous effects data, i.e., the data in which the median amino-acid effect was removed.

We estimated the relative contributions of the experimental error and the effect of synonymous mutations in the data by comparing the impact of *replicate, codon* and *medium* (i.e., whether salt was added or not) using the following ANOVA model with data between replicates 2 and 3 of both the standard and the high-salinity environment for 30C:

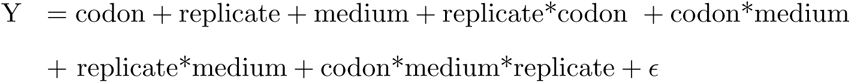

where Y corresponds to the normalized selection coefficient, *codon* to a fixed factor corresponding to the 64 codons present in the data, *replicate* to a fixed factor pertaining to the arbitrary replicate number 2 or 3 for each environment, *medium* is a fixed factor corresponding to the presence or absence of high salt concentration in the medium and *ϵ* corresponds to the residual error. Additionally, we estimated effect size by calculating *η*^2^ (i.e., the ratio of the variance explained by a predictor to the total variance explained by the entire model - (Levine and Hullett, 2002)) for each of the model terms, using the etasq function of the *R* package sjstats (Liidecke, 2017). To assess the variability of our estimates, we performed the analysis for 1000 posterior samples. We find that the fitness effects of codon changes contribute more to the total variance of the model than variation in replicates, indicating that we can detect overall effects of codon changes, despite the presence of experimental error Fig. S7.

#### Quantifying the effect size of synonymous mutations

To quantify the effect size of synonymous codon changes, we performed a linear regression for each amino-acid (including all amino-acids with 3 or more codons) and calculated *η*^2^ for the *codon* term as proxy for effect size (Levine and Hullett, 2002). The regression per amino-acid was performed within each environment and took into account *residue solvent accessibility* (i.e., whether the position was buried or exposed). Pooling of positions was performed to allow for the testing of codon effect within an amino-acid. To minimize potential differences arising from pooling positions, we separated the data into buried and exposed positions according to solvent accessibility of the residue. Additionally, using an ANOVA model we tested how the estimated effect size per amino-acid (using *η*^2^ as dependent variable) varied across *environment* and *amino-acid*.

We performed three different but related types of analyses to quantify the average fitness effects of synonymous mutations. Firstly, we focused solely on the 15 mutations that are synonymous to the reference sequence (similar to Bank *et al.*, 2014). Here, we computed the medians of the maximum and minimum effect size, and the standard deviation from 1000 samples of the posterior. Secondly, across the whole data set, we computed the descriptive statistics of the differences between each codon and the average amino-acid effect of this codon. Finally, we compared the distributions of the absolute pairwise differences between amino-acid effects, synonymous-codon effects, and samples from the posterior of the same codon. For the environments 30N and 30S (30^°^C with normal and high salinity) we performed all analyses across the available 3 and 2 replicates, respectively, and confirmed that our conclusions remain qualitatively similar (results not shown).

### 2.5 Potential mechanisms underlying the effect of synonymous mutations on fitness

There are several mechanisms through which synonymous mutations can affect protein translation (reviewed in Plotkin and Kudla, 2011). In this study we focused on whether codon usage frequency or predicted mRNA stability (using RNA melting temperature as a proxy) can predict effects of synonymous mutations(Presnyak *et al.*, 2015).

Firstly, to enable the inclusion of codon frequency patterns in yeast into our regression models, we obtained the relative abundance of each codon in the yeast genome from the Codon Usage Database (http://www.kazusa.or.jp/codon/cgi-bin/showcodon.cgi?species=4932).

Secondly, synonymous mutations may affect translation through different stability of the mRNA generated by different codons. To obtain predictions of how mRNA stability is affected by synonymous mutations, we used the prediction software mfold (Zuker *et al.*, 1999; Markham and Zuker, 2008), for 25^°^C, 30^°^C and 36^°^C and with high salt concentrations (0.5 M Na^+^), with physiological concentrations of salt (0.015 M Na^+^), and 0.001 M Mg^2+^, respectively. As input, we used sequences spanning 135 nucleotides of the Hsp90 protein in yeast. To obtain these sequences, we added 54 nucleotides flanking both 5’ and 3’ sides of the region of interest (complete sequences were obtained from https://www.addgene.org/41188/sequences/). From each of these data sets, we selected the conformation with the highest melting temperature (Tm), as highest-stability reference point.

Since Hsp90 is a chaperone involved in the response to thermal stress as well as in the regulation of osmotic stress (Yang *et al.*, 2006; Boucher *et al.*, 2014), we tested which factors can explain variation in codon fitness effects. For that we performed model selection using the leaps package (Lumley, 2017). We started with the full model with all factors *(Temperature, Medium, Codon frequency, Melting temperature, Residue position* and *GC content*) and their interactions and proceeded by backward selection. We selected the best models under three different criteria - BIC (Bayesian Information Criteria), adjusted R^2^ and Mallow’s C_*p*_. To select the best model we calculated the credibility interval, based on 1000 posterior samples, for AIC, BIC, R^2^ and adjusted R^2^ and performed model comparison with an ANOVA analyses.

### 2.6 Effect of synonymous mutations on the topography and the dynamics of adaptive walks in codon fitness landscapes

To quantify the impact of effects of synonymous mutations coding for the same amino-acid on the topography of the fitness landscape, we compared the single-effect landscape with the averaged landscape. For the single-effect landscapes (Fig. 1A3) the effect of each codon was directly obtained from the experimental data. For the averaged landscape (Fig. 1A2) we assigned to every codon that coded for the same amino-acid the same pooled amino-acid estimate obtained from *empiricIST.*

Each amino-acid position in our data set corresponds to a complete multi-allelic fitness landscape with 4^3^ = 64 genotypes. We characterized the prevalence of epistasis in the resulting 9.6 = 54 fitness landscapes using several fitness landscape statistics. We estimated 1) the roughness-to-slope ratio (Aita *et al.*, 2001; Szendro *et al.*, 2013; Bank *et al.*, 2016) to quantify the relative deviations from an additive model; 2) the multi-allelic gamma statistics (Bank *et al.*, 2016; Ferretti *et al.*, 2016) to characterize the prevalence and type of epistasis in the landscape. Additionally, to test the impact of synonymous mutations on the evolutionary dynamics on the landscape we estimated: 1) the number of local peaks (Szendro *et al.*, 2013); and 2) the length and variance in the length of potential adaptive walks in the landscapes (Neidhart and Krug, 2011; Szendro *et al.*, 2013). Probabilities of adaptive walks were computed analytically under the strong-selection weak-mutation approximation, following Bank *et al.* (2016). Credibility of the estimates was assessed by computing the fitness landscape statistics for 100 posterior samples. Differences between averaged and single-effect landscapes were assessed by comparing the lower and upper 2.5% boundary of the credibility estimates. Differences were considered significant if the lower credibility interval from the averaged landscape and the upper credibility interval of the single-effect landscape (and vice-versa) did not overlap for each of the statistics. We did not perform multiple testing adjustment for this analysis.

All analyses were performed with R (R version 3.3.3) (R Core Team, 2017) or Mathematica 11 (version 11.2) (Wolfram Research, Inc., 2017). The complete documentation of all analyses, which allows for the reiteration of all steps, is available from the Dryad Digital Repository https://doi.org/10.5061/dryad.k7jm5hp

### 3 Results & Discussion

#### 3.1 The distribution of fitness effects of synonymous mutations

Previous studies have shown that synonymous mutations can directly affect fitness (e.g. Lind *et al.*, 2010; Firnberg *et al.*, 2014; Hunt *et al.*, 2014) and impact the ability of populations to adapt to new environments (Bailey *et al.*, 2014; Agashe *et al.*, 2016). For example, Bailey *et al.* (2014) found that two synonymous mutations were responsible for adaptation of *Pseudomonas fluorescens* to a new medium by increasing the expression of a gene involved in glucose metabolism. In a more recent study in *Methylobacterium extorquens*, Agashe *et al.* (2016) found that the deleterious effect of synonymous mutations in a medium with methylamine as the sole carbon source could be rescued by different mutations, including four synonymous mutations that increased transcription and protein production levels. The impact of synonymous mutations at the genome-wide level was also found in patterns of codon usage bias (synonymous codons are used at different frequencies) across genomes. Evidence from studies within and between species support the role of direct selection on synonymous sites in various genes (DuMont *et al.*, 2004; Singh *et al.*, 2007; Hershberg and Petrov, 2009; Ran and Higgs, 2010; Shah and Gilchrist, 2011; Choi and Aquadro, 2016; Sun *et al.*, 2016). A first piece of evidence for synonymous effects in the studied region of Hsp90 came from Bank *et al.* (2014), who reported that one of the 15 mutations synonymous to the parental sequence had a significantly deleterious effect in 4 out of 6 environments (Fig. 9 in Bank *et al.*, 2014). In order to quantify the distribution of synonymous fitness across different amino-acid backgrounds, we applied *empiricIST* to the data set from Bank *et al.* (2014), which provided us with the growth rate of all 576 possible codon mutations across a 9 amino-acid region of Hsp90 in *Saccharomyces cerevisiae* in 6 different experimental conditions, estimated from bulk competitions. We extracted the synonymous fitness contribution of each mutation by subtracting the mean amino-acid effect.

##### 3.1.1 The effect size of synonymous mutations

Although most of the 15 mutations that are synonymous to the wild type are of similar effect, some individual synonymous mutations present as much as 1% of fitness change (Codon AAC, Fig S8). Since the data set includes all 64 codon mutations for each residue position, we obtained a larger set of synonymous mutations by extracting the effect of synonymous mutations across all amino acids. By default, these mutations include the effect of amino-acid changes in relation to the wild type plus possible effects of synonymous mutations. To eliminate the amino-acid effect, for all mutations we subtracted the estimated average effect of the corresponding amino acid (see Methods). As a result of this, the DFE of the synonymous mutations is concentrated around 0. Its shape is clearly different from the DFE of the non-synonymous mutations (Fig S9), with much lower effect sizes. As observed for the synonymous mutations to the wild type, the average effect size of synonymous mutations varies across environments, from 0.001 in 30S to 0.004 in 36N Table S3B). However, effect sizes are also highly variable (Fig S9) and can reach up to 0.019 in 25N, 0.010 in 25S, 0.022 in 30N, 0.014 in 30S, 0.045 in 36N and 0.023 in 36S (Table S3B)

To quantify the effect of synonymous mutations in comparison with the effect of non-synonymous mutations and experimental error, we calculated the absolute pairwise differences between 1000 random pairs of amino-acids, codons, posterior samples and replicates (Table S4). This allowed us to estimate the average effect of: 1) an amino-acid change (non-synonymous mutations), 2) a codon change within the same amino acid (synonymous mutations), 3) variation between posterior samples (estimation error). Expectedly, absolute pairwise differences between mutants that changed the amino acid have a much stronger effect than synonymous mutations (Table S4, see Material & Methods). Overall differences between synonymous mutations are higher than two random draws of the posterior. On average, the effects of synonymous mutations are larger in the 36N environment (Table S4, Fig. S10), where Hsp90 is expected to be more important for organism survival (Yang *et al.*, 2006; Boucher *et al.*, 2014; Mishra *et al.*, 2016).

Overall, our assessment shows that the size of synonymous fitness effects is non-negligible but only slightly above the experimental detection limit. Therefore, we restrict ourselves to statistical arguments about the distribution of synonymous effects in this study rather than identifying specific mutations which would likely result in a high false discovery rate.

##### 3.1.2 The beneficial tail of the distribution of synonymous fitness effects

The distribution of fitness effects contains information about the availability of beneficial mutations (Orr, 2005, 2010). It is of particular interest to study the shape of the beneficial tail of this distribution as it determines various aspects regarding the nature of adaptive walks (Orr, 2010; Eyre-Walker, 2006). Using the same data as in the present study, Bank *et al.* (2014) previously found that for all environments except 25S, the beneficial tail of the full distribution of fitness effects most likely belonged to the Weibull domain. This suggested that populations were close to a well-defined optimum, and the available beneficial mutations would be of similar and small size (Orr, 2010; Joyce *et al.*, 2008; Bank *et al.*, 2014).

We used the tail shape estimator from *empiricIST* to estimate the tail shape of the distribution of beneficial synonymous mutations (i.e., the distribution of the beneficial contributions to the amino acid effects). We find that the shape parameter of the fitted Generalized Pareto Distribution is most likely positive in all environments, which indicates that the resulting shape of the beneficial tail belongs to the Fréchet domain (Fig. 3) (Orr, 2010; Joyce *et al.*, 2008). Distributions from this domain are characterized by many mutations of small effect, along with few mutations of large and unpredictable effect (Joyce *et al.*, 2008; Neidhart and Krug, 2011; Jain and Seetharaman, 2011). Such a shape of the distribution of synonymous effects makes sense both intuitively and with respect to the reported examples of large-effect synonymous mutations (Bailey *et al.*, 2014; Agashe *et al.*, 2016): a majority of synonymous mutations may not affect fitness at all, whereas specific ones could indeed significantly affect fitness. This would also explain why synonymous effects are seldomly detected, as the rarity of large effects implies that a large number of mutations has to be screened to obtain a positive result.

**Figure 3:**
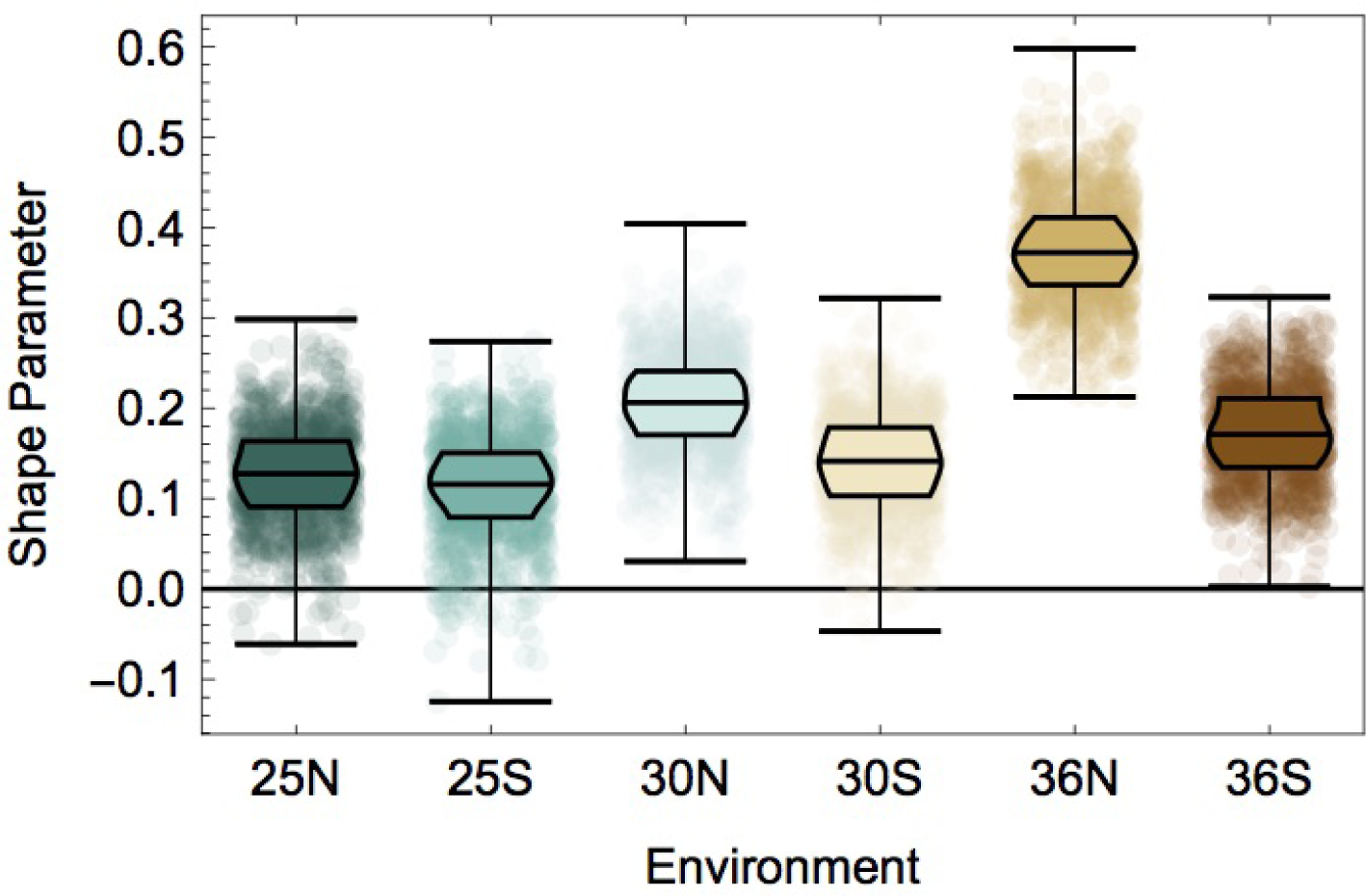
Shape parameter estimates of the beneficial tail of the distribution of synonymous effects. In all environments, the estimated shape parameter of the tail is positive, which indicates that the distribution of synonymous effects belongs to the Frechet domain (i.e., that it can be characterized by a heavy-tailed distribution). This implies the presence of many nearly-neutral and few larger-effect mutations. The shape parameter was estimated using the tail shape estimator from *empiricIST* using the information of 1000 samples from the posterior distribution. Environmental conditions are indicated as the combination of temperature (25C, 30C and 36C) and salinity (N = normal and S = high salinity).

##### 3.1.3 Relationship between observed synonymous effects and potential underlying biological mechanisms

Synonymous mutations can affect fitness by altering speed and accuracy of translation, and mRNA folding and stability (Drummond and Wilke, 2008; Kudla *et al.*, 2009; Zhou *et al.*, 2009; Sharp *et al.*, 2010; Plotkin and Kudla, 2011; Shabalina *et al.*, 2013; Presnyak *et al.*, 2015; Yu *et al.*, 2015; Knöppel *et al.*, 2016; Brule and Grayhack, 2017). It has been proposed that protein folding may be affected more significantly by changes in translation accuracy for buried (structural) positions, as they are often involved in the formation of crucial secondary and tertiary structures of the protein (Drummond and Wilke, 2008; Zhou *et al.*, 2009; Saunders and Deane, 2010). The usage of different synonymous codons could therefore allow cells to slow down or arrest protein production in response to sudden environmental changes and to optimize resource production (Zhang *et al.*, 2009; Fredrick and Ibba, 2010; Tuller *et al.*, 2010). We evaluated whether the effects of synonymous mutations that we observe can be explained by variation in codon preference or mRNA stability. To this end, we analyzed a full linear model incorporating temperature, medium composition, residue position, melting temperature of mRNAs, GC content, and codon usage frequency, as well as all possible interactions of those factors.

No clear predictors of codon fitness emerged from this analysis. The best model indicated that fitness effects of synonymous mutations are affected by interactions between residue positions and temperature, medium composition, mRNA melting temperature, GC content, and codon usage frequency Table S5); however, only 1.4% of the variance in fitness effects could be attributed to this combination of factors. There are various reasons that could explain this inconclusive result. Firstly, the synonymous effect sizes could be too small compared with the experimental uncertainty to yield a clear result. This problem should be amplified by the observed shape of the distribution of synonymous effects; if only few mutations have an effect, the statistical power to detect this effect in the full data set will be very low. Secondly, we considered diverse amino-acid positions and environments. Intuitively, it seems plausible that at each of the positions, different biological mechanisms could contribute to synonymous fitness effects. Thirdly, our analysis is based on a distribution of synonymous fitness effects that was observed on top of an amino-acid effect in a conserved region of the protein, which could blur the true distribution of synonymous effects. Thus, larger data sets based on synonymous mutations to a common reference will be necessary for a better statistical assessment of the factors underlying the distribution of synonymous fitness effects.

#### 3.2 The shape of the codon fitness landscape with and without synonymous effects

Having established that there is a non-negligible distribution of synonymous fitness effects, it is natural to ask how considering such effects changes a given fitness landscape. In the following section, we analyze the 54 64-genotype fitness landscapes of single amino-acid positions that are contained in our data set. In contrast to the section above, we now also consider the amino-acid effects of mutations and compare the shape of the fitness landscape when (1) all codons for the same amino acid are assigned the same effect (averaged landscape) and when (2) all codons have individually estimated effects (single-effect landscape).

##### 3.2.1 Epistasis in the codon fitness landscape

We investigated the effect of synonymous mutations on the topography of the fitness landscape by comparing the prevalence and type of epistasis for averaged and single-effect landscapes (see Fig. 1 A3, 1 A2, Material & Methods) for each of the 9 amino-acid positions across 6 environments. For all 54 landscapes, we computed two statistics: the roughness-to-slope ratio *r/s* (Szendro *et al.*, 2013) and the locus-specific gamma statistic (Ferretti *et al.*, 2016). The roughness-to-slope ratio quantifies the prevalence of epistasis by comparing the deviation of the landscape from an additive model with the magnitude of the fitness effects (Carneiro and Hartl, 2010; Schenk *et al.*, 2013). The γ_*i*→*j*_ statistic measures the correlation of fitness effects of the same mutations in a single-step distance across all genetic backgrounds. Whereas the roughness-to-slope ratio describes the landscape by means of only a single value, γ_*i*→*j*_ results in a representation of the landscape by means of 2*L* values, where *L* is the number of loci. This epistatic footprint makes heterogeneity of epistasis in the landscape visible, and can thus indicate epistatic signals at the level of single loci (e.g. Bank *et al.*, 2016).

The roughness-to-slope-ratio indicates that all but one of the codon landscapes are highly epistatic (*r/s >* 1), with the magnitude of the roughness-to-slope ratio varying across amino-acid positions and environments (Fig. S11). Single-effect landscapes tend to be more epistatic (larger roughness-to-slope ratio) than averaged landscapes, although this difference is in general small. Interestingly, in few cases the single-effect landscape has a smaller roughness-to-slope ratio than its corresponding averaged landscape. This is noteworthy because in this case the consideration of synonymous fitness effects makes the landscape less rugged/more linear, which is opposite to the intuitive expectation that adding variation in synonymous effects should increase the number of peaks and thus the prevalence of epistasis in the landscape. At high salinity, the roughness-to-slope ratio tends to be larger than in normal environments, and also the difference in the roughness-to-slope ratios between amino-acid positions and between averaged and single-effect landscapes is larger (Fig. S11). The stronger epistatic signal observed in the high-salinity environments could be caused by the combination of low absolute growth rates observed in high salinity conditions that result in larger relative fitness differences of the mutations (c.f. Table 1 in Bank *et al.*, 2014), and larger experimental uncertainty (Fig. S6) in this environment. This indicates that one needs to be cautious when interpreting roughness-to-slope ratios across data sets, because the measure may be confounded by experimental differences rather than genuine changes in the epistatic component of the landscape.

Computing the γ_*i*→*j*_ statistic per codon position confirms that averaged and single-effect landscapes tend to display a similar strength of epistasis within amino-acid position and environment on a global scale (Fig. 4, Fig. S12). Only when γ_*i*→*j*_ is computed for individual pairs of nucleotide substitutions, larger differences in epistasis appear across the resulting epistatic profiles (see Fig. S13). The γ_*i*→*j*_ statistic per codon position shows smaller differences between environments than the roughness-to-slope ratio. As the gamma statistic is based on the correlation and not the effect size of fitness effects across genetic backgrounds, it is less sensitive to differences in mutational effect sizes and experimental error. The largest systematic differences in the codon-position-wise strength of epistasis are found when comparing the order in which mutations occur. Gamma measures obtained from the epistatic effect of non-synonymous mutations (γ_1→2_, γ_2→1_) in general display strong epistasis (Fig. 4), compared to gamma measures obtained from (mostly) synonymous mutations (γ_1→3_, γ_2→3_, γ_3→1_, γ_3→2_). Thus, the structure of the codon table (i.e., the existence of synonymous and non-synonymous mutations) leaves a distinctive signal in all fitness landscapes, but the signal looks similar for averaged and single-effect landscapes. Splitting this signal into its components caused by individual pairs of nucleotides illustrates extensive local heterogeneity of epistasis across codon fitness landscapes and between single-effect and averaged landscapes (see sup Fig. S13) and indicates the potential for different dynamics of adaptive walks, which is discussed in the next Section. However, this measure describes each codon fitness landscape by a set of 216 values, which makes it difficult to obtain quantitative comparisons. Nevertheless, Fig. S13 shows qualitatively that every codon fitness landscape has indeed a different epistatic profile. This local heterogeneity of the fitness landscapes is not well captured by averaging summary statistics such as the roughness-to-slope ratio and the codon-position-based gamma statistic.

**Figure 4:**
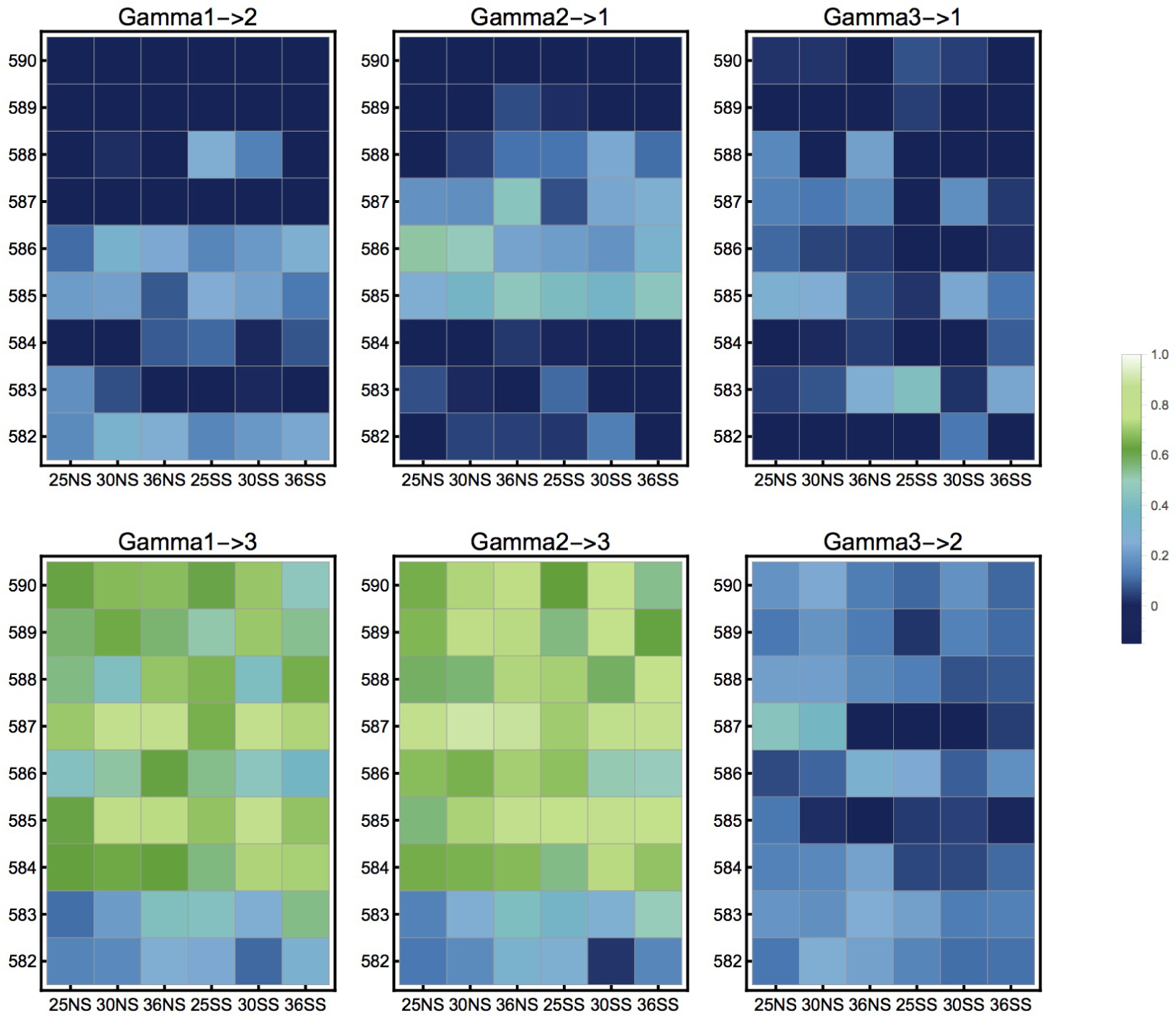
Gamma statistics of pairs of codon positions for single-effect landscapes across amino-acid positions (*y* axis) and environments *x* axis). In general, interactions of non-synonymous mutations (γ_1→2_, γ_1→2_) are more epistatic, than non-synonymous mutations in the background of synonymous mutations. There is no systematic variation across environments (*x* axis), but there seems to be a systematic impact of amino-acid position on the strength of epistasis (*y* axis). Specifically, position 582 shows qualitatively stronger epistasis for both γ_1→3_ and γ_2→3_ across all environments. This indicates that synonymous mutations may play a relatively larger role at constraining evolutionary paths at this position.

#### 3.3 Impact of synonymous mutations on adaptive walks

Including synonymous mutations changes the topography of the landscape, which may affect the accessibility of different mutational paths by creating additional peaks and sinks in the fitness landscape. To quantify the impact of synonymous effects on adaptive walks, we calculated the number of optima, the mean expected length of adaptive walks, and the variance in the number of steps for the single-effect and averaged landscapes. We based our calculation on the assumption of the strong-selection weak-mutation limit (Gillespie, 1984), in which evolution happens by means of sequential beneficial substitutions that result in an adaptive walk that ends in a fitness peak (e.g. Orr, 2005; Schoustra *et al.*, 2009; Frank, 2014; Zagorski *et al.*, 2016). We define a fitness peak as any genotype with fitness higher than all single-step mutational neighbors. For averaged landscapes, in which all synonymous mutations are assigned equal fitness, we consider a fitness plateau spanned by synonymous codons as a single local optimum if all non-synonymous codons in a distance of a single nucleotide step have lower fitness (as in Fig. 1 A3).

By definition, the number of fitness peaks in the averaged landscape has to be lower or equal to that of the single-effect landscape. Indeed, we find that there is a large difference in the number of fitness peaks between single-effect and averaged landscapes (Fig. 5, Fig. S14). This difference is environment-dependent and also varies across amino-acid positions (Fig. 5, Fig. S14), and it is accompanied by a larger between-environment variation in the number of peaks in the single-effect landscapes. For most environments and positions, averaged landscapes have only 1 or 2 fitness peaks (Fig. 5, Fig. S14). Conversely, among the single-effect landscapes, 25^°^C stands out with a consistently large number of peaks across all amino-acid positions (mean across positions for single-effect: 5.568, mean across positions for averaged landscape: 2.593). The much larger number of fitness peaks in the single-effect landscape suggests that evolution on the ‘true’ fitness landscape that includes effects of synonymous mutations is less predictable (Lobkovsky *et al.*, 2011; De Visser and Krug, 2014).

**Figure 5:**
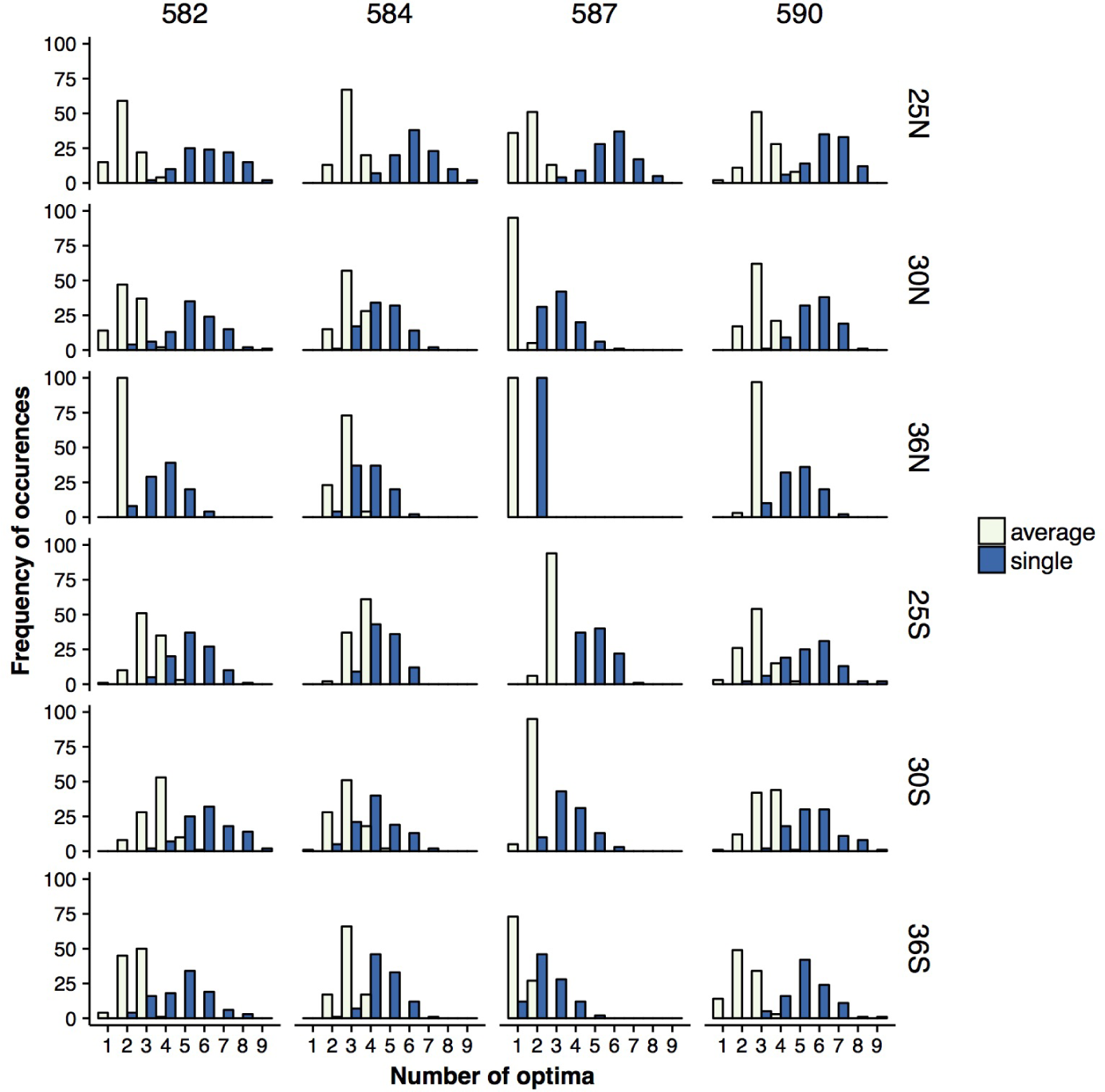
Figure 5: Number of optima observed from 100 posterior samples of single-effect (dark blue) and averaged (light yellow) landscapes for positions 582, 584, 586 and 590 (from left to right) across environments. (See Fig. S14 for the complete set of loci.) The number of optima is always larger in single-effect than in averaged landscapes. The number of optima is smaller at high temperatures, which may indicate increased constraints to adaptation. The large difference between the number of peaks in averaged and single landscapes suggests that synonymous mutations can affect adaptation to a new environment by trapping the population at a local optimum. The mean and median of the distributions are significantly different according to Welch Two Sample t-test and Wilcoxon test, *p* < 0.00001 after Bonferroni correction for 54 comparisons.

As synonymous mutations are expected to have a stronger effect in buried amino-acid positions (Drummond and Wilke, 2008), differences between adaptive walks on single-effect and averaged landscapes should be larger in buried positions. However, we do not see consistent variation between the two landscape types between buried or exposed positions (see Material & Methods), which suggests that the impact of synonymous mutations is not solely due to effects on protein folding, i.e. not strongly correlated with the solvent accessibility of the residues.

Biologically, a larger difference in predicted adaptive walks between averaged and single-effect landscapes points to a greater importance of synonymous fitness effects. The pronounced differences that we observed at cold temperature could stem both from the smaller absolute growth rate of the wild type in this environment, which results in larger relative effects of mutations (i.e., small-effect mutations could become more visible), and from a reduced need for functional Hsp90 at cold temperature (i.e., Hsp90 is not so necessary), which could result in a larger number of (synonymous) adaptive solutions that are connected to the fine-tuning of the protein. In support of this hypothesis, we observe fewer optima and longer and more variable adaptive walks in the single-effect landscapes at 36N (Table. S6, Table. S7), which is in agreement with the importance of Hsp90 at high temperatures (Hietpas *et al.*, 2013; Bank *et al.*, 2014; Boucher *et al.*, 2014; Mishra *et al.*, 2016), which may leave only few options for improvement. This is consistent with the small proportion of beneficial mutations across the whole DFE observed by Bank *et al.* (2014) in this condition.

Our results allow for an interesting thought experiment regarding the impact of synonymous mutations on evolution across populations of different sizes. Our results add to the notion that synonymous fitness effects exist but are small on average. According to the nearly-neutral theory, such small fitness effects will only be visible to selection if the population size is large (Ohta, 1992). When they become visible in large populations, synonymous fitness effects create additional peaks in the organism’s fitness landscape, in which adaptation can become stalled.

In such a situation, bottlenecks (i.e., sudden drops in the population size), which can occur under natural scenarios and are also frequently imposed in experiments, may render synonymous mutations effectively neutral. This erases the previous fitness peak and allows the population to continue their adaptive walk on an averaged-type fitness landscape. Thus, by opening mutational paths and erasing synonymous fitness peaks, a (temporally) smaller population size could speed up adaptation and increase its predictability (Wright, 1931; Jain *et al.*, 2011). Caused by the different effect size and distribution of non-synonymous versus synonymous mutations, this effect is in contrast to the slowdown of adaptation and decrease of predictability of evolution in small populations proposed in standard population-genetic theory (Orr, 2000; Lanfear *et al.*, 2014).

## 4 Conclusion

The impact of the codon table on the evolutionary dynamics on fitness landscapes has received little attention. This is a consequence of the vast size of the nucleotide space and the resulting dimensionality of the fitness landscape, which has led to most studies restricting themselves to the amino-acid level. Using selection coefficient estimates obtained with *empiricIST*, a new software for the estimation of growth rates from deep mutational scanning data, we characterized the distribution of synonymous fitness effects and investigated the consequences of including synonymous mutations when characterizing the fitness landscape of single amino-acid positions across environments. Interestingly, we found support for a heavy-tailed distribution of beneficial synonymous effects across all environments, suggestive of a distribution of fitness effects with many small-or-no effect mutations and few mutations of potentially large effects. This is in line with the current population-genetics literature, in which the importance of accounting for synonymous fitness effects is discussed controversially. We demonstrate that synonymous mutations can impact the topography of the fitness landscape and affect adaptation in an environment-dependent fashion. Importantly, we show that synonymous fitness effects can directly impact both the path and endpoint of an adaptive walk by creating additional fitness peaks. This highlights the importance of their consideration in the study of fitness landscapes.

## 5 Acknowledgments

We thank the members of the Bank lab for discussion of the manuscript. We thank Daniel Bolon and Ryan Hietpas for originally obtaining the data analyzed in this publication. This work was supported by Fundagao Calouste Gulbenkian and an ERC Starting Grant to JDJ. I. Fragata was supported by a postdoctoral fellowship from FCT (Fundagao para a Ciencia e a Tecnologia) within the project JPIAMR/0001/2016.

## References

Agashe D, Martinez-Gomez NC, Drummond DA, Marx CJ (2013). Good codons, bad transcript: Large reductions in gene expression and fitness arising from synonymous mutations in a key enzyme. Molecular Biology and Evolution 30: 549–560.

Agashe D, Sane M, Phalnikar K, Diwan GD, Habibullah A, Martinez-Gomez NC, et al. (2016). Large-Effect Beneficial Synonymous Mutations Mediate Rapid and Parallel Adaptation in a Bacterium. Molecular Biology and Evolution 33: 1542–1553.

Aita T, Iwakura M, Husimi Y (2001). A cross-section of the fitness landscape of dihydrofolate reductase. Protein Engineering, Design and Selection 14: 633–638.

Bailey SF, Hinz A, Kassen R (2014). Adaptive synonymous mutations in an experimentally evolved Pseudomonas fluorescens population. Nature Communications 5: 1–7.

Bali V, Bebok Z (2015). Decoding mechanisms by which silent codon changes influence protein biogenesis and function. International Journal of Biochemistry and Cell Biology 64: 58–74.

Bank C, Hietpas RT, Wong A, Bolon DN, Jensen JD (2014). A Bayesian MCMC approach to assess the complete distribution of fitness effects of new mutations: Uncovering the potential for adaptive walks in challenging environments. Genetics 196: 841–852.

Bank C, Matuszewski S, Hietpas RT, Jensen JD (2016). On the (un)predictability of a large intragenic fitness landscape. Proceedings of the National Academy of Sciences 113: 14085–14090.

Bataillon T, Zhang T, Kassen R (2011). Cost of adaptation and fitness effects of beneficial mutations in Pseudomonas fluorescens. Genetics 189: 939–49.

Beisel CJ, Rokyta DR, Wichman HA, Joyce P (2007). Testing the extreme value domain of attraction for distributions of beneficial fitness effects. Genetics 176: 2441–2449.

Boone EL, Merrick JR, Krachey MJ (2014). A Hellinger distance approach to MCMC diagnostics. Journal of Statistical Computation and Simulation 84: 833–849.

Boucher JI, Cote P, Flynn J, Jiang L, Laban A, Mishra P, et al. (2014). Viewing protein fitness landscapes through a next-gen lens. Genetics 198: 461–471.

Brule CE, Grayhack EJ (2017). Synonymous Codons: Choose Wisely for Expression. Trends in Genetics 33: 283–297.

Carneiro M, Hartl DL (2010). Adaptive landscapes and protein evolution. Proceedings of the National Academy of Sciences 107: 1747–1751.

Chevin LM (2011). On measuring selection in experimental evolution. Biology letters 7: 210–3.

Choi JY, Aquadro CF (2016). Recent and Long-Term Selection Across Synonymous Sites in Drosophila ananassae. Journal of Molecular Evolution 83: 50–60.

De Visser JAGM, Krug J (2014). Empirical fitness landscapes and the predictability of evolution. Nature Reviews Genetics 15: 480–490.

Drummond DA, Wilke CO (2008). Mistranslation-Induced Protein Misfolding as a Dominant Constraint on Coding-Sequence Evolution. Cell 134: 341–352.

DuMont VB, Fay JC, Calabrese PP, Aquadro CF (2004). DNA variability and divergence at the Notch locus in Drosophila melanogaster and D. simulans: A case of accelerated synonymous site divergence. Genetics 167: 171–185.

Eyre-Walker A (2006). The genomic rate of adaptive evolution. Trends in Ecology and Evolution 21: 569–575.

Ferretti L, Schmiegelt B, Weinreich D, Yamauchi A, Kobayashi Y, Tajima F, et al. (2016). Measuring epistasis in fitness landscapes: The correlation of fitness effects of mutations. Journal of Theoretical Biology 396: 132–143.

Firnberg E, Labonte JW, Gray JJ, Ostermeier M (2014). A comprehensive, high-resolution map of a Gene’s fitness landscape. Molecular Biology and Evolution 31: 1581–1592.

Fowler DM, Fields S (2014). Deep mutational scanning: A new style of protein science. Nature Methods 11: 801–807.

Frank SA (2014). Generative models versus underlying symmetries to explain biological pattern. Journal of Evolutionary Biology 27: 1172–1178.

Fredrick K, Ibba M (2010). How the sequence of a gene can tune its translation. Cell 141: 227–229.

Gelman A, Roberts G, Gilks W (1996). Efficient metropolis jumping rules. In: Bayesian statistics, Oxford Science Publications, vol. 5, pp. 599–607.

Gillespie JH (1983). A simple stochastic gene substitution model. Theoretical Population Biology 23: 202–215.

Gillespie JH (1984). Molecular Evolution Over the Mutational Landscape. Evolution 38: 1116–1129.

Gorter FA, Aarts MGM, Zwaan BJ, de Visser JAGM (2018). Local Fitness Landscapes Predict Yeast Evolutionary Dynamics in Directionally Changing Environments. Genetics 1: 307–322.

Hershberg R, Petrov DA (2009). General rules for optimal codon choice. PLoS Genetics 5: 1–10.

Hietpas R, Roscoe B, Jiang L, Bolon DN (2012). Fitness analyses of all possible point mutations for regions of genes in yeast. Nature Protocols 7: 1382–1396.

Hietpas RT, Bank C, Jensen JD, Bolon DNA (2013). Shifting fitness landscapes in response to altered environments. Evolution 67: 3512–3522.

Hietpas RT, Jensen JD, Bolon DNA (2011). Experimental illumination of a fitness landscape. Proceedings of the National Academy of Sciences 108: 7896–7901.

Hunt RC, Simhadri VL, Iandoli M, Sauna ZE, Kimchi-Sarfaty C (2014). Exposing synonymous mutations. Trends in Genetics 30: 308–321.

Jain K, Krug J, Park SC (2011). Evolutionary advantage of small populations on complex fitness landscapes. Evolution 65: 1945–1955.

Jain K, Seetharaman S (2011). Multiple adaptive substitutions during evolution in novel environments. Genetics 189: 1029–1043.

Joyce P, Rokyta DR, Beisel CJ, Orr HA (2008). A general extreme value theory model for the adaptation of DNA sequences under strong selection and weak mutation. Genetics 180: 1627–1643.

Knöppel A, Näsvall J, Andersson DI (2016). Compensating the Fitness Costs of Synonymous Mutations. Molecular Biology and Evolution 33: 1461–1477.

Kudla G, Murray AW, Tollervey D, Plotkin JB (2009). Coding-Sequence Determinants of Gene Expression in Escherichia coli. Science 324: 255–258.

Lanfear R, Kokko H, Eyre-Walker A (2014). Population size and the rate of evolution. Trends in Ecology and Evolution 29: 33–41.

Levine TR, Hullett CR (2002). Eta Squared, Partial Eta Squared, and Misreporting of Effect Size in Communication Research. Human Communication Research 28: 612–625.

Lind PA, Berg OG, Andersson DI (2010). Mutational robustness of ribosomal protein genes. Science 330: 825–827.

Lobkovsky AE, Wolf YI, Koonin EV (2011). Predictability of evolutionary trajectories in fitness landscapes. PLoS Computational Biology 7: 1–11.

Lumley T (2017). leaps: Regression subset Selection. R package version 3.0.

Lüdecke D (2017). sjstats: Statistical Functions for Regression Models. R package version 0.14.0. URL: https://CRAN.R-project.org/package=sjstats

Markham NR, Zuker M (2008). UNAFold: Software for nucleic acid folding and hybridization. Methods in Molecular Biology 453: 3–31.

Matuszewski S, Hildebrandt ME, Ghenu AH, Jensen JD, Bank C (2016). A statistical guide to the design of deep mutational scanning experiments. Genetics 204: 77–87.

Mishra P, Flynn JM, Starr TN, Bolon DN (2016). Systematic Mutant Analyses Elucidate General and Client-Specific Aspects of Hsp90 Function. Cell Reports 15: 588–598.

Neidhart J, Krug J (2011). Adaptive walks and extreme value theory. Physical Review Letters 107: 1–4.

O’Brien EP, Ciryam P, Vendruscolo M, Dobson CM (2014). Understanding the influence of codon translation rates on cotranslational protein folding. Accounts of Chemical Research 47: 1536–1544.

Ohta T (1992). The nearly neutral theory of molecular evolution. Annual Review of Ecology and Systematics 23: 263–286.

Orr HA (2000). The rate of adaptation in asexuals. Genetics 155: 961–968.

Orr HA (2005). The genetic theory of adaptation: A brief history. Nature Reviews Genetics 6: 119–127.

Orr HA (2010). The population genetics of beneficial mutations. Philosophical Transactions of the Royal Society B: Biological Sciences 365: 1195–1201.

Parmley JL, Hurst LD (2007). How do synonymous mutations affect fitness? BioEssays 29: 515–519.

Perfeito L, Ghozzi S, Berg J, Schnetz K, Lässig M (2011). Nonlinear Fitness Landscape of a Molecular Pathway. PLoS Genetics 7: e1002160.

Plotkin JB, Kudla G (2011). Synonymous but not the same: The causes and consequences of codon bias. Nature Reviews Genetics 12: 32–42.

Presnyak V, Alhusaini N, Chen YH, Martin S, Morris N, Kline N, et al. (2015). Codon optimality is a major determinant of mRNA stability. Cell 160: 1111–1124.

R Core Team (2017). R: A Language and Environment for Statistical Computing. R Foundation for Statistical Computing, Vienna, Austria. URL: https://www.R-project.org/

Ran W, Higgs PG (2010). The influence of anticodon-codon interactions and modified bases on codon usage bias in bacteria. Molecular Biology and Evolution 27: 2129–2140.

Roberts GO, Rosenthal JS, et al. (2001). Optimal scaling for various metropolis-hastings algorithms. Statistical science 16: 351–367.

Rodriguez-Verdugo A, Carrillo-Cisneros D, Gonzalez-Gonzalez A, Gaut BS, Bennett AF (2014). Different tradeoffs result from alternate genetic adaptations to a common environment. Proceedings of the National Academy of Sciences 111: 12121–12126.

Rokyta DR, Beisel CJ, Joyce P, Ferris MT, Burch CL, Wichman HA (2008). Beneficial Fitness Effects Are Not Exponential for Two Viruses. Journal of molecular evolution 67: 368–376.

Sauna ZE, Kimchi-Sarfaty C (2011). Understanding the contribution of synonymous mutations to human disease. Nature Reviews Genetics 12: 683–691.

Saunders R, Deane CM (2010). Synonymous codon usage influences the local protein structure observed. Nucleic Acids Research 38: 6719–6728.

Schenk MF, Szendro IG, Salverda MLM, Krug J, De Visser JAGM (2013). Patterns of epistasis between beneficial mutations in an antibiotic resistance gene. Molecular Biology and Evolution 30: 1779–1787.

Schoustra SE, Bataillon T, Gifford DR, Kassen R (2009). The properties of adaptive walks in evolving populations of fungus. PLoS Biology 7: 1–10.

Shabalina SA, Spiridonov NA, Kashina A (2013). Sounds of silence: Synonymous nucleotides as a key to biological regulation and complexity. Nucleic Acids Research 41: 2073–2094.

Shah P, Gilchrist MA (2011). Explaining complex codon usage patterns with selection for translational efficiency, mutation bias, and genetic drift. Proceedings of the National Academy of Sciences 108: 10231–10236.

Sharp PM, Emery LR, Zeng K (2010). Forces that influence the evolution of codon bias. Philosophical Transactions of the Royal Society B: Biological Sciences 365: 1203–1212.

Singh ND, Bauer DuMont VL, Hubisz MJ, Nielsen R, Aquadro CF (2007). Patterns of mutation and selection at synonymous sites in Drosophila. Molecular Biology and Evolution 24: 2687–2697.

Sun Y, Tamarit D, Andersson SG (2016). Switches in Genomic GC Content Drive Shifts of Optimal Codons under Sustained Selection on Synonymous Sites. Genome Biology and Evolution p. evw201.

Szendro IG, Schenk MF, Franke J, Krug J, De Visser JAGM (2013). Quantitative analyses of empirical fitness landscapes. Journal of Statistical Mechanics Theory and Experiment p. P01005.

Tuller T, Waldman YY, Kupiec M, Ruppin E (2010). Translation efficiency is determined by both codon bias and folding energy. Proceedings of the National Academy of Sciences 107: 3645–3650.

Weinreich DM, Delaney NF, DePristo MA, Hartl DL (2006). Darwinian evolution can follow only very few mutational paths to fitter proteins. Science 312: 111–114.

Wenger JW, Piotrowski J, Nagarajan S, Chiotti K, Sherlock G, Rosenzweig F (2011). Hunger artists: Yeast adapted to carbon limitation show trade-offs under carbon sufficiency. PLoS Genetics 7: 1–17.

Wolfram Research, Inc (2017). Mathematica v. 11.2. Champaign, Illinois, USA.

Wright S (1931). Evolution in mendelian populations. Genetics 16: 97–159.

Wu NC, Dai L, Olson CA, Lloyd-Smith JO, Sun R (2016). Adaptation in protein fitness landscapes is facilitated by indirect paths. eLife 5: 1–21.

Yang XX, Maurer KCT, Molanus M, Mager WH, Siderius M, Van Der Vies SM (2006). The molecular chaperone Hsp90 is required for high osmotic stress response in Saccharomyces cerevisiae. FEMS Yeast Research 6: 195–204.

Yu CH, Dang Y, Zhou Z, Wu C, Zhao F, Sachs MS, et al. (2015). Codon Usage Influences the Local Rate of Translation Elongation to Regulate Co-translational Protein Folding. Molecular Cell 59: 744–754.

Zagorski M, Burda Z, Waclaw B (2016). Beyond the Hypercube: Evolutionary Accessibility of Fitness Landscapes with Realistic Mutational Networks. PLoS Computational Biology 12: 1–18.

Zhang G, Hubalewska M, Ignatova Z (2009). Transient ribosomal attenuation coordinates protein synthesis and co-translational folding. Nature Structural and Molecular Biology 16: 274–280.

Zhou T, Weems M, Wilke CO (2009). Translationally optimal codons associate with structurally sensitive sites in proteins. Molecular Biology and Evolution 26: 1571–1580.

Zuker M, Mathews D, Turner D (1999). Algorithms and Thermodynamics for RNA secondary structure prediction: a pratical guide. RNA biochemistry and biotechnology pp. 1–33.

